# Goldilocks calcium and the mitochondrial respiratory chain: too much, too little, just right

**DOI:** 10.1101/2022.04.12.488015

**Authors:** Eloisa A. Vilas-Boas, João Victor Cabral-Costa, Vitor M. Ramos, Camille C. Caldeira da Silva, Alicia J. Kowaltowski

## Abstract

Calcium (Ca^2+^) is a key regulator in diverse intracellular signaling pathways, and has long been implicated in metabolic control and mitochondrial function. Mitochondria can actively take up large amounts of Ca^2+^, thereby acting as important intracellular Ca^2+^ buffers and affecting cytosolic Ca^2+^ transients. Excessive mitochondrial matrix Ca^2+^ is known to be deleterious due to opening of the mitochondrial permeability transition pore (mPTP) and consequent membrane potential dissipation, leading to mitochondrial swelling, rupture, and cell death. Moderate Ca^2+^ within the organelle, on the other hand, can directly or indirectly activate mitochondrial matrix enzymes, possibly impacting on ATP production. Here, we aimed to determine in a quantitative manner if extra or intramitochondrial Ca^2+^ modulate oxidative phosphorylation in mouse liver mitochondria and intact hepatocyte cell lines. To do so, we monitored the effects of more modest versus supra-physiological increases in cytosolic and mitochondrial Ca^2+^ on oxygen consumption rates. Isolated mitochondria present increased respiratory control ratios (a measure of oxidative phosphorylation efficiency) when incubated with low (2.4 ± 0.6 μM) and medium (22.0 ± 2.4 μM) Ca^2+^ concentrations in the presence of complex I-linked substrates pyruvate plus malate and α-ketoglutarate, respectively, but not complex II-linked succinate. In intact cells, both low and high cytosolic Ca^2+^ led to decreased respiratory rates, while ideal rates were present under physiological conditions. High Ca^2+^ decreased mitochondrial respiration in a substrate-dependent manner, mediated by mPTP. Overall, our results uncover a Goldilocks effect of Ca^2+^ on liver mitochondria, with specific “just right” concentrations that activate oxidative phosphorylation.

## 1. Introduction

Calcium (Ca^2+^) is an important second messenger and participates in a myriad of cellular functions. Mitochondria are one of the central intracellular regulators of Ca^2+^ homeostasis (1), as they can actively take up the ion down their electrochemical potential (2,3). While the affinity for Ca^2+^ uptake is lower than for the endoplasmic reticulum (ER), mitochondria can absorb very large amounts of the ion from the cytosol (4–6). Mitochondria can also release Ca^2+^, and are thus recognized as important rheostats for cytosolic Ca^2+^ dynamics and signaling. Furthermore, mitochondria also participate in crosstalk signals between intracellular compartments, such as with the ER, through microdomains formed between both organelles in which ions have distinct concentrations (7).

Ca^2+^ uptake across the inner mitochondrial membrane into the mitochondrial matrix occurs through the mitochondrial Ca^2+^ uniporter complex (MCU) (8–10). The membrane potential generated by the electron transport chain (ETC), with a negative matrix charge, provides the electrochemical force necessary for positively charged ions, such as Ca^2+^, to enter. Ca^2+^ efflux from the mitochondrial matrix occurs through the Na^+^/Ca^2+^ exchanger (NCLX) (11) and by a recently identified Ca^2+^/H^+^ exchanger (12,13), located in the inner mitochondrial membrane (IMM).

Mitochondrial Ca^2+^ transients are believed to provide a link between cytosolic Ca^2+^ signaling and the control of cellular energy demand by regulating ATP production. Pyruvate dehydrogenase, isocitrate dehydrogenase, and α-ketoglutarate dehydrogenase within the mitochondrial matrix have increased activities in the presence of Ca^2+^, as uncovered through *in vitro* studies (14–17). For these activities to translate into enhanced oxidative phosphorylation *in vivo,*two conditions need to be met: first, the change in enzymatic activity promoted by Ca^2+^ needs to be sufficient to overcome rate-limiting steps in oxidative phosphorylation and, second, Ca^2+^ concentrations need to be below those that hamper mitochondrial function. Mitochondrial Ca^2+^ overload leads to swelling of the organelle and opening of the mitochondrial permeability transition pore (mPTP), often culminating in loss of function and cell death (18–20). This process has been implicated in various diseases (21,22).

In theory, and when present at lower concentrations than those that lead to mPTP opening, Ca^2+^ should activate mitochondrial electron transport and ATP production. Previous studies have explored the modulation of isolated mitochondrial enzyme activities by Ca^2+^ (14,23), showing increases in substrate affinity. Other studies using primary hepatocytes exposed to vasopressin and glucagon have unveiled an important role of increased cytosolic Ca^2+^ on mitochondrial Ca^2+^ levels, which result in the regulation of mitochondrial matrix volume (24,25) and the activation of matrix dehydrogenases (26,27). However, the effects of different Ca^2+^ concentrations on overall mitochondrial metabolic fluxes, with emphasis on mitochondrial oxygen consumption rates, have not been extensively determined.

We explore this knowledge gap here, and determine if extra or intramitochondrial Ca^2+^ modulates oxygen consumption fluxes under controlled side-by-side comparison conditions using different substrates, respiratory states, and their quantitative responses to specific, calibrated, and distinct amounts of Ca^2+^. We used mitochondria isolated from the liver, since a rich metabolic regulation by Ca^2+^ occurs in this tissue. As different Ca^2+^amounts can be present in distinct isolated mitochondrial preparations, in order to minimize possible confounding results, we performed daily calibrations, which allowed us to uncover activating Ca^2+^ effects with more accuracy, as these were often subtle and concentration-specific. We also explore how mitochondrial Ca^2+^ transport influences oxidative phosphorylation in intact hepatocyte-derived cells. Our results demonstrate that Ca^2+^ concentrations greatly impact mitochondrial respiration, with a Goldilocks-type effect, in which both too much and too little Ca^2+^ limit oxidative phosphorylation, but the “just right” concentration promotes significant activation.

## 2. Results

### 2.1. Ca^2+^ increases liver oxidative phosphorylation efficiency in a substrate- and concentration-dependent manner

To gain insight into the effects of extramitochondrial Ca^2+^ additions on mitochondrial respiration, we used isolated mouse liver mitochondria in the presence of different substrates and measured oxygen consumption over time using an Oroboros high-resolution oxygraph. Ca^2+^ was added at different concentrations, followed by additions of ADP (to induce oxidative phosphorylation, state 3), oligomycin (oligo, to inhibit ATP synthase and measure respiration dependent on the proton leak, state 4), and CCCP (to induce maximum electron transport). We determined residual Ca^2+^ concentrations present after mitochondrial isolation daily, and then adjusted the amount of Ca^2+^ added so it was equal between biological replicates. Based on the amount of Ca^2+^ added and excess EGTA present in the buffer, we calculated the amount of free Ca^2+^ to which mitochondria were exposed, classifying conditions as no added Ca^2+^, low (2.4 ± 0.6 μM), medium (22.0 ± 2.4 μM), and high (52.9 ± 2.5 μM) Ca^2+^ concentrations.

Complex I substrates such as pyruvate plus malate were chosen initially to study the effects of Ca^2+^ on electron transport and oxidative phosphorylation, due to the known enhancement of pyruvate dehydrogenase (PDH) affinity in response to Ca^2+^ signaling (14–17). Ca^2+^ uptake traces with low, medium, and high Ca^2+^ concentrations are shown in Fig. 1A. The downward deflection in the curve within 2-3 min after the addition of extramitochondrial Ca^2+^ indicates a decrease in extramitochondrial Ca^2+^ measured with the fluorophore Ca^2+^ Green, due to uptake by mitochondria. High Ca^2+^ led to mitochondrial dysfunction with these substrates, as indicated by the inability to maintain the Ca^2+^ uptake over time (Fig. 1A), and was thus not included in further experiments. O2 consumption traces were then measured using low and medium Ca^2+^ additions, and are shown as representative oxygen consumption rates (OCR) in Figs. 1B-D and quantified results in panels E-H.

**Figure 1.**
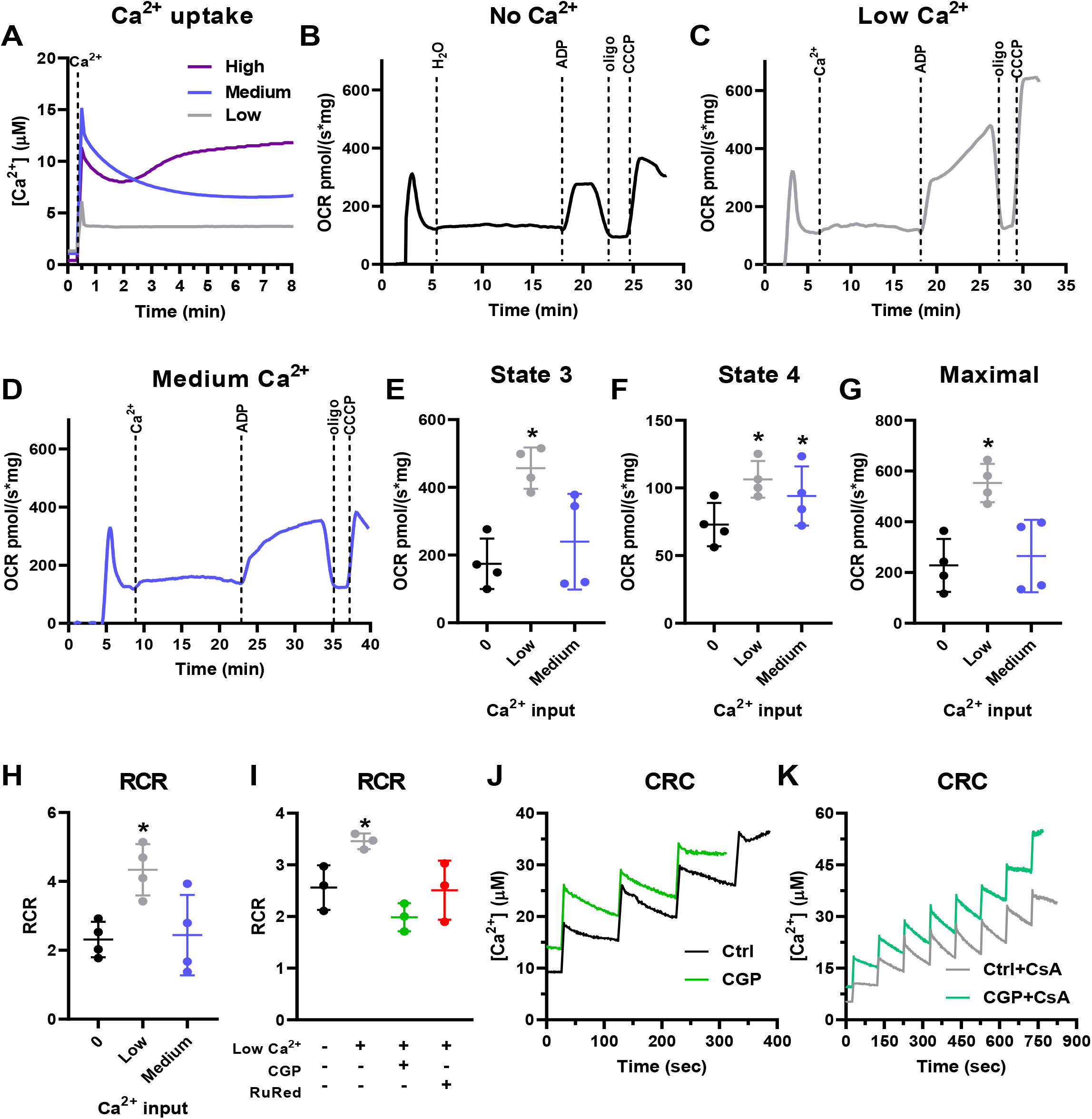
Ca^2+^ uptake increases liver mitochondrial oxidative phosphorylation supported by pyruvate plus malate. Isolated mouse liver mitochondria (600 μg) were incubated with 1 mM pyruvate + 1 mM malate. **A)** Representative Ca^2+^ uptake curves using Ca^2+^ Green 5N to measure extramitochondrial Ca^2+^ in the presence of low Ca^2+^ (2.4 ± 0.6 μM, gray line), medium Ca^2+^ (22.0 ± 2.4 μM, blue line), and high Ca^2+^ (52.9 ± 2.5 μM, purple line) concentrations. **B-D)** Representative O2 consumption rate (OCR) traces using a high-resolution Oxygraph-2k (O2k) respirometer in response to zero Ca^2+^ **(B)**, low Ca^2+^ **(C)**, and medium Ca^2+^ **(D)** additions, followed by injection of 1 mM ADP, 1 μM oligomycin (oligo), and 0.5 μM CCCP. **E)** State 3, after ADP addition. **F)** State 4, after oligomycin addition. **G)** Maximal respiration after CCCP addition. **H)** Respiratory control ratios (RCR, state 3 / state 4). **I)** RCR in the presence of low Ca^2+^ and 10 μM CGP-37157 (CGP) or 10 μM ruthenium red (RuRed). n = 4 independent experiments. *p<0.05 versus control using one-way ANOVA followed by Šidak. **J)** Ca^2+^ retention capacity (CRC) under control conditions (Ctrl) and in the presence of CGP. **K)** CRC with cyclosporin A (CsA) in the absence or presence of CGP.

We found that the presence of low Ca^2+^ concentrations strongly increased state 3 respiration, when oxidative phosphorylation is active (Fig. 1E), and state 3u, after the uncoupler CCCP (Fig. 1G). State 4 respiration, which occurs in the absence of ATP synthase activity and is limited by the inner membrane impermeability to protons, was stimulated by both low and medium Ca^2+^ concentrations when supported by pyruvate plus malate (Fig. 1F). Respiratory control ratios (RCR, Fig. 1H) are the ratio between state 3 and state 4 oxygen consumption, and used as a proxy for oxidative phosphorylation efficiency, as they show the relative stimulation of oxygen consumption by production of mitochondrial ATP (28). We found that RCRs were increased at low, but not medium, Ca^2+^ concentrations (Fig. 1H). This shows that oxidative phosphorylation supported by pyruvate plus malate has a tight relationship with mitochondrial Ca^2+^, and is more effective within a very specific range.

We investigated next if the increase in oxidative phosphorylation efficiency observed was dependent on mitochondrial Ca^2+^ uptake or cycling, using pharmacological inhibitors of the MCU (Ruthenium red, RuRed), which prevents Ca^2+^ uptake, and of the NCLX (CGP-37157, CGP), which prevents Ca^2+^ extrusion from mitochondria (Fig. 1I). Both RuRed and CGP reversed the increase in RCR promoted by low Ca^2+^concentrations; this indicates that Ca^2+^ must enter the mitochondrial matrix to exert this effect, as RuRed prevents this uptake. The effect of CGP could also indicate that increased RCRs are dependent on Ca^2+^ cycling, as it inhibits Ca^2+^ extrusion from mitochondria through Ca^2+^ exchange with Na^+^ (11). Alternatively, CGP treatment of mitochondria, by promoting more Ca^2+^ retention in the matrix, could induce mPTP opening and thus decrease RCRs. To investigate this possibility, we measured Ca^2+^ retention capacity using pyruvate plus malate with CGP in the presence or absence of the mPTP inhibitor cyclosporin A (CsA). We found that the addition of CGP decreases Ca^2+^ retention capacity to 11.8 μM, in comparison with 24.5 μM under control conditions (Fig. 1J). Additionally, incubation with CsA, as expected, greatly amplified Ca^2+^ retention capacity (43.9 μM with CsA under control conditions and 32.4 μM in the presence of CGP plus CsA) (Fig. 1K). CsA also decreased the effect of CGP (Fig. 1K), suggesting that the decrease in RCR promoted by NCLX inhibition by CGP is related to the induction of mPTP due to higher intramitochondrial Ca^2+^ accumulation.

To further investigate the role of Ca^2+^ on oxidative phosphorylation, we performed similar experiments using α-ketoglutarate (Fig. 2), another substrate metabolized by a dehydrogenase in which affinity is modulated by Ca^2+^*in vitro* (14–17). Once again, only low and medium, but not high Ca^2+^ concentrations, could be taken up and retained by mitochondria respiring on α-ketoglutarate (Fig. 2A). Using this substrate, medium Ca^2+^concentrations (22.0 ± 2.4 μM Ca^2+^) increased states 3 and the RCR, without changes in maximal respiration (Fig. 2B-H). Low Ca^2+^ concentrations did not present a significant effect when α-ketoglutarate was used as the substrate. The beneficial effect of Ca^2+^ on mitochondrial oxidative phosphorylation supported by α-ketoglutarate was partially attenuated by inhibition of Ca^2+^ release by the NCLX using CGP, and totally prevented by inhibition of Ca^2+^ entry with RuRed (Fig. 2I), demonstrating again the need for Ca^2+^entry into the matrix for the stimulatory effect. These results show that the specific “sweet spot” for enhanced electron transport capacity in mitochondria induced by Ca^2+^ varies with the substrate used. When using α-ketoglutarate as substrate, the Ca^2+^ retention capacity was smaller than when using pyruvate plus malate (both in the absence and presence of CGP) (Fig. 2J): 7.2 μM under control conditions and 6.7 μM with CGP. CsA again increased all CRCs (Fig. 2K): 28.4 μM with CsA and 22.5 μM with CsA plus CGP.

**Figure 2.**
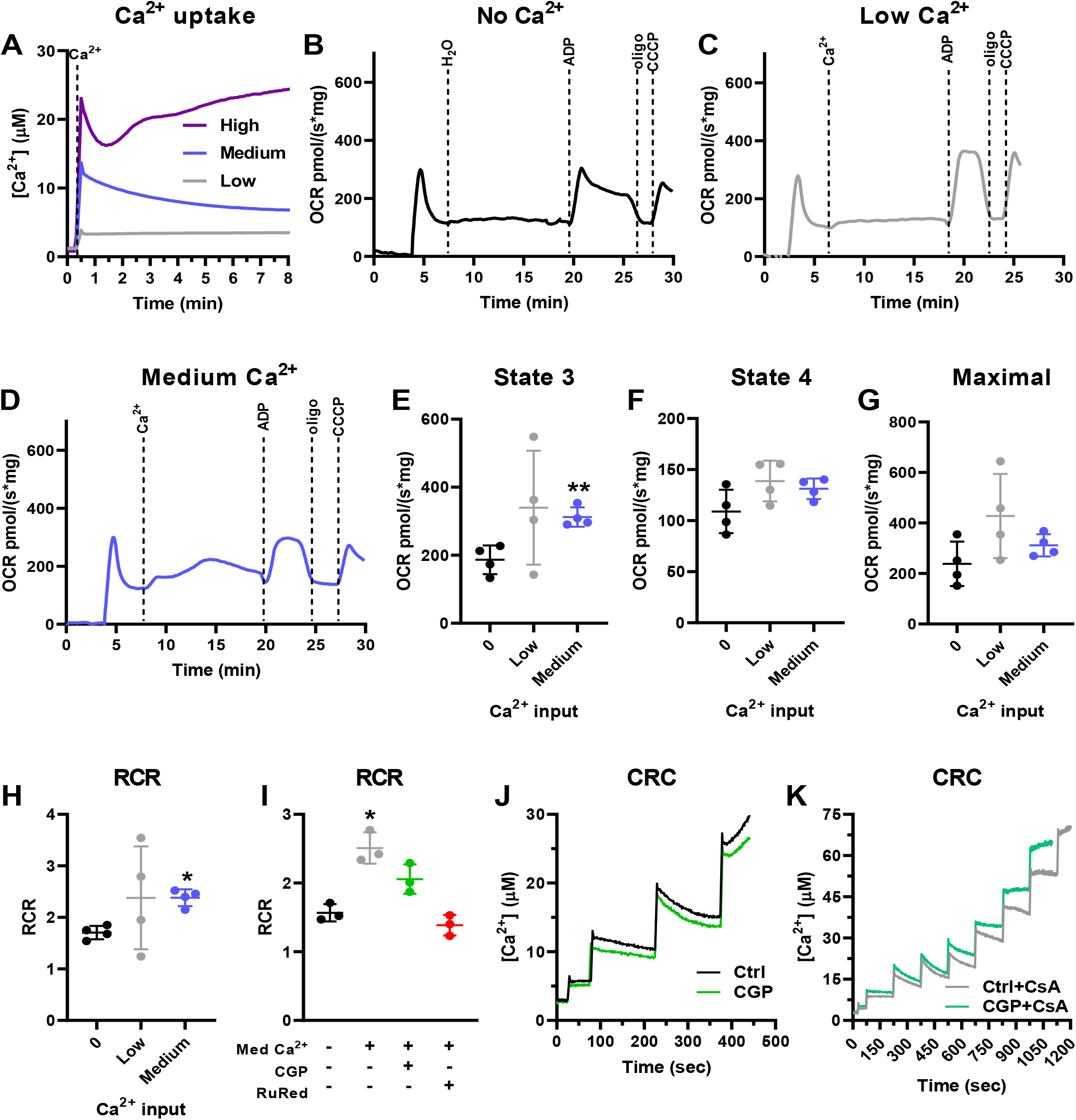
Ca^2+^ uptake increases liver mitochondrial oxidative phosphorylation supported by α-ketoglutarate. Isolated mouse liver mitochondria (600 μg) were energized with 1 mM α-ketoglutarate. **A)** Representative Ca^2+^ uptake curves using Ca^2+^ Green 5N to measure extramitochondrial Ca^2+^ in the presence of low Ca^2+^ (2.4 ± 0.6 μM, gray line), medium Ca^2+^ (22.0 ± 2.4 μM, blue line), and high Ca^2+^ (52.9 ± 2.5 μM, purple line). **B-D)** Representative OCR traces with zero Ca^2+^ **(B)**, low Ca^2+^ **(C)**, and medium Ca^2+^ **(D)**, followed by injection of 1 mM ADP, 1 μM oligomycin (oligo), and 0.5 μM CCCP. **E)** State 3, after ADP addition. **F)** State 4, after oligomycin addition. **G)** Maximal respiration, after CCCP addition. **H)** Respiratory control ratios (RCR, state 3 / state 4). **I)** RCR in the presence of medium Ca^2+^ and CGP or RuRed. n = 4 independent experiments; *p<0.05 and **p<0.01 versus control using one-way ANOVA followed by Šidak. **J)** Ca^2+^ retention capacity (CRC) under control conditions (Ctrl) and in the presence of CGP. **K)** CRC with cyclosporin A (CsA) in the absence or presence of CGP.

To further investigate substrate-specific effects of Ca^2+^, we performed similar experiments using the complex II substrate succinate in the presence of complex I inhibition with rotenone, to ensure no effect of contaminating or downstream NADH-reducing substrates. With succinate, mitochondria were able to retain a wider range of Ca^2+^ loads (Fig. 3A), so low, medium, and high concentrations were tested for their respiratory effects. Interestingly, using succinate, none of these Ca^2+^ concentrations led to a significant increase in state 3, maximal respiration or RCR (Fig. 3F, H, I). A small increase in state 4, in which respiration is not linked to oxidative phosphorylation, was observed with the medium Ca^2+^ concentration (Fig. 3G), suggesting enhanced proton leak, possibly due to Ca^2+^ cycling or mPTP induction in a subpopulation of the mitochondrial suspension.

**Figure 3.**
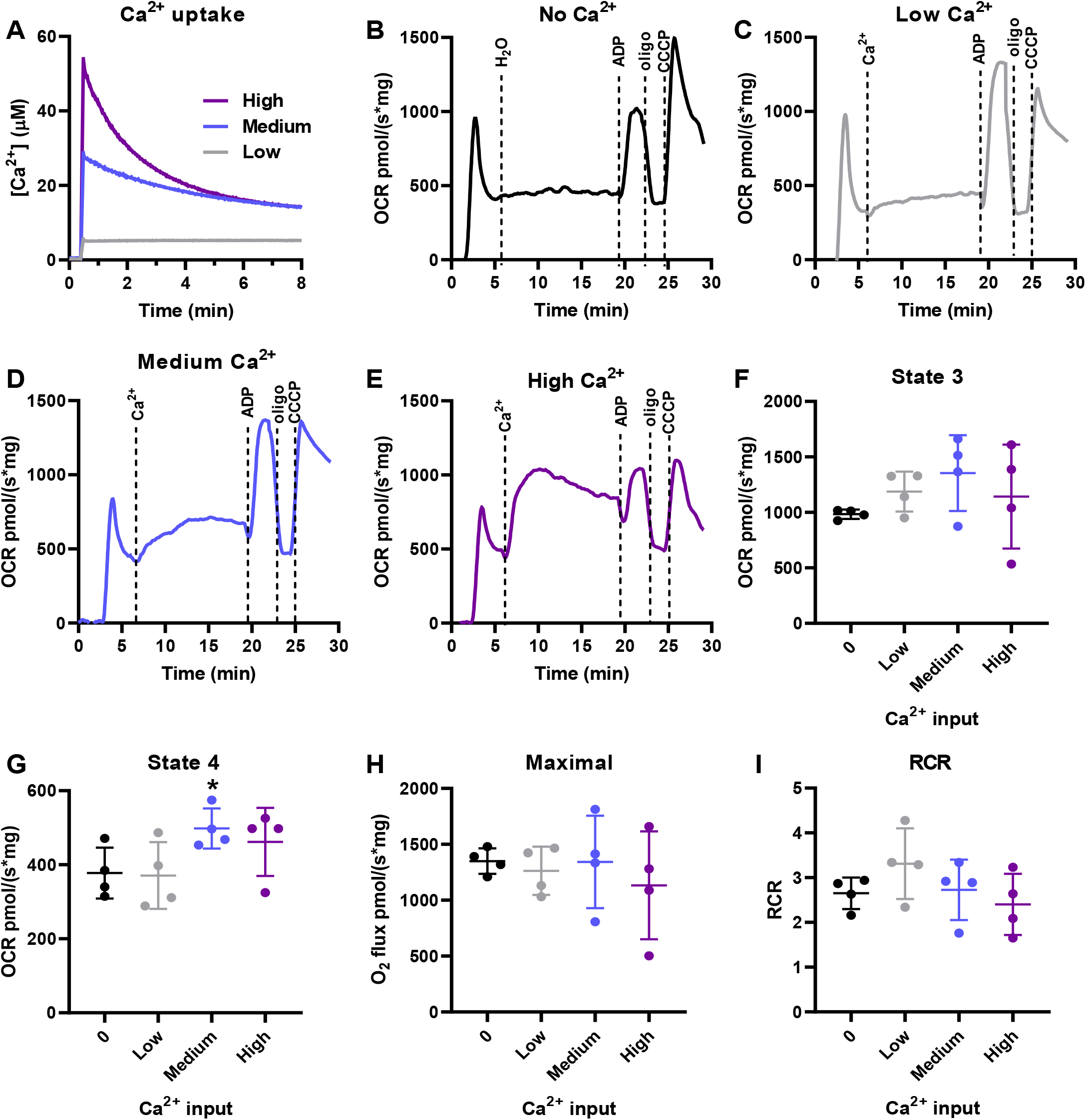
Ca^2+^ uptake does not increase liver mitochondrial oxidative phosphorylation supported by succinate. Isolated mouse liver mitochondria (200 μg) were energized with 1 mM succinate in the presence of 1 μM rotenone. **A)** Representative Ca^2+^ uptake curves using Ca^2+^ Green 5N to measure extramitochondrial Ca^2+^ in the presence of low Ca^2+^ (2.4 ± 0.6 μM, gray line), medium Ca^2+^ (22.0 ± 2.4 μM, blue line), and high Ca^2+^ (52.9 ± 2.5 μM, purple line). **B-E)** Representative OCR traces in response to zero Ca^2+^ **(B)**, low Ca^2+^ **(C)**, medium Ca^2+^**(D)**, and high Ca^2+^ **(E)**, followed by injection of 1 mM ADP, 1 μM oligomycin (oligo), and 0.5 μM CCCP. **F)** State 3, after ADP addition. **G)** State 4, after oligomycin addition. **H)** Maximal respiration after CCCP addition. **I)** Respiratory control ratios (RCR, state 3 / state 4). n = 4 independent experiments; *p<0.01 versus control using one-way ANOVA followed by Šidak.

Since we measured an increase in oxidative phosphorylation efficiency by Ca^2+^with two substrates metabolized by matrix NADH-linked dehydrogenases, we questioned whether Ca^2+^ influenced the formation of NAD(P)H. NAD(P)H autofluorescence was measured with pyruvate plus malate in the absence and presence of low Ca^2+^ concentrations (Fig. 4A), and with α-KG with and without medium Ca^2+^ concentrations (Fig. 4B). We saw no changes promoted by Ca^2+^ in NAD(P)H quantities under basal conditions, nor in the presence of any of the respiratory modulators used. We then wondered if the effects of Ca^2+^ could involve changes in mitochondrial membrane potentials (ΔΨm), and measured ΔΨm under our experimental conditions (Fig. 4C,E, typical traces, quantified in panels D,F). ΔΨm using pyruvate plus malate as substrates is sustained (with or without Ca^2+^) throughout the measurements (Fig. 4C), with no changes between control conditions (no Ca^2+^) and low Ca^2+^ in the different respiratory states analyzed (Fig. 4D): basal state, after 10 minutes of Ca^2+^ addition, state 3 (after ADP), state 4 (after oligomycin), and state 3u (after CCCP). When using α-KG as the substrate, we noticed a gradual loss of ΔΨm with time (Fig. 4E). This could be due to mPTP formation, possibly in mitochondrial subpopulations, since we do not see significant Ca^2+^ efflux under these conditions. Despite this, ΔΨm quantifications were not significantly affected by Ca^2+^ (Fig. 4F), indicating changes in protonmotive force cannot be mechanistically ascribed to the Ca^2+^ effects on electron transport.

**Figure 4.**
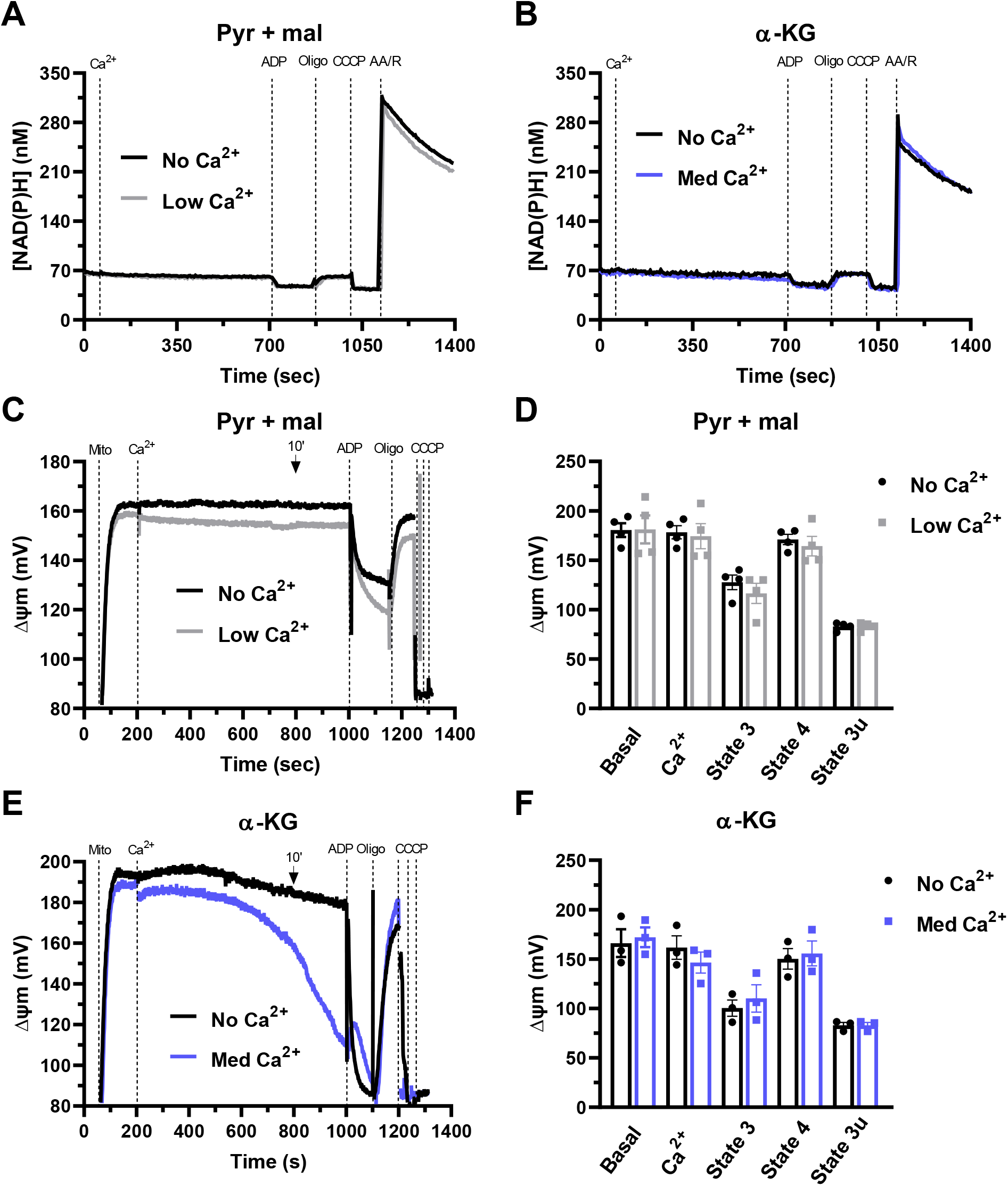
NAD(P)H levels and mitochondrial membrane potentials (ΔΨm) are unaltered by respiratory-activating Ca^2+^ concentrations. Isolated mouse liver mitochondria (600 μg) were energized with 1 mM pyruvate + 1 mM malate (pyr + mal) or 1 mM α-ketoglutarate (α-KG). Representative NAD(P)H (**A-B)** and ΔΨm **(C,E)** time scans using pyr + mal with zero Ca^2+^ (no Ca^2+^, black line) or low Ca^2+^ (2.4 ± 0.6 μM, gray line) **(A, C)**, or α-KG with zero Ca^2+^ (no Ca^2+^, black line) or medium Ca^2+^ (22.0 ± 2.4 μM, blue line) **(B, E)**. Fluorescence was assessed after the addition of Ca^2+^, followed by 1 mM ADP, 1 μM oligomycin (oligo), 0.5 μM CCCP, and 1 μM rotenone + 1 μM antimycin A (R/AA). Values are expressed as nM **(A-B)** or mV **(C-F)**. **D-F)** Quantification of ΔΨm with pyr + mal **(D)** or α-KG **(F)** under basal conditions, after 10 minutes of Ca^2+^ addition, in state 3 (after ADP), state 4 (after oligomycin), and state 3u (after CCCP). Results are expressed as means ± SEM (B, C, E, F) of 3–4 independent experiments, no significant changes were detected using one-way ANOVA followed by Šidak.

Taken together, results using isolated mouse liver mitochondria and various substrates show that a low increase in mitochondrial Ca^2+^ enhances oxidative phosphorylation capacity and efficiency. This positive effect depends on the substrate used and requires Ca^2+^ entry into the mitochondrial matrix.

### 2.2. Both low and high cytosolic Ca^2+^ impair mitochondrial respiration in cultured hepatocyte-derived cells

We investigated next how mitochondrial and cytosolic Ca^2+^ modulate electron transport and oxidative phosphorylation in intact cells (Fig. 5). Since primary isolated hepatocytes undergo major metabolic alterations during and after isolation (29), defeating the purpose of using a freshly isolated cell, we used AML12 cells, a non-tumoral hepatocyte cell line (30), and thus a more physiological liver cell model. OCRs were measured in plated cells using a Seahorse Extracellular Flux system (31).

**Figure 5.**
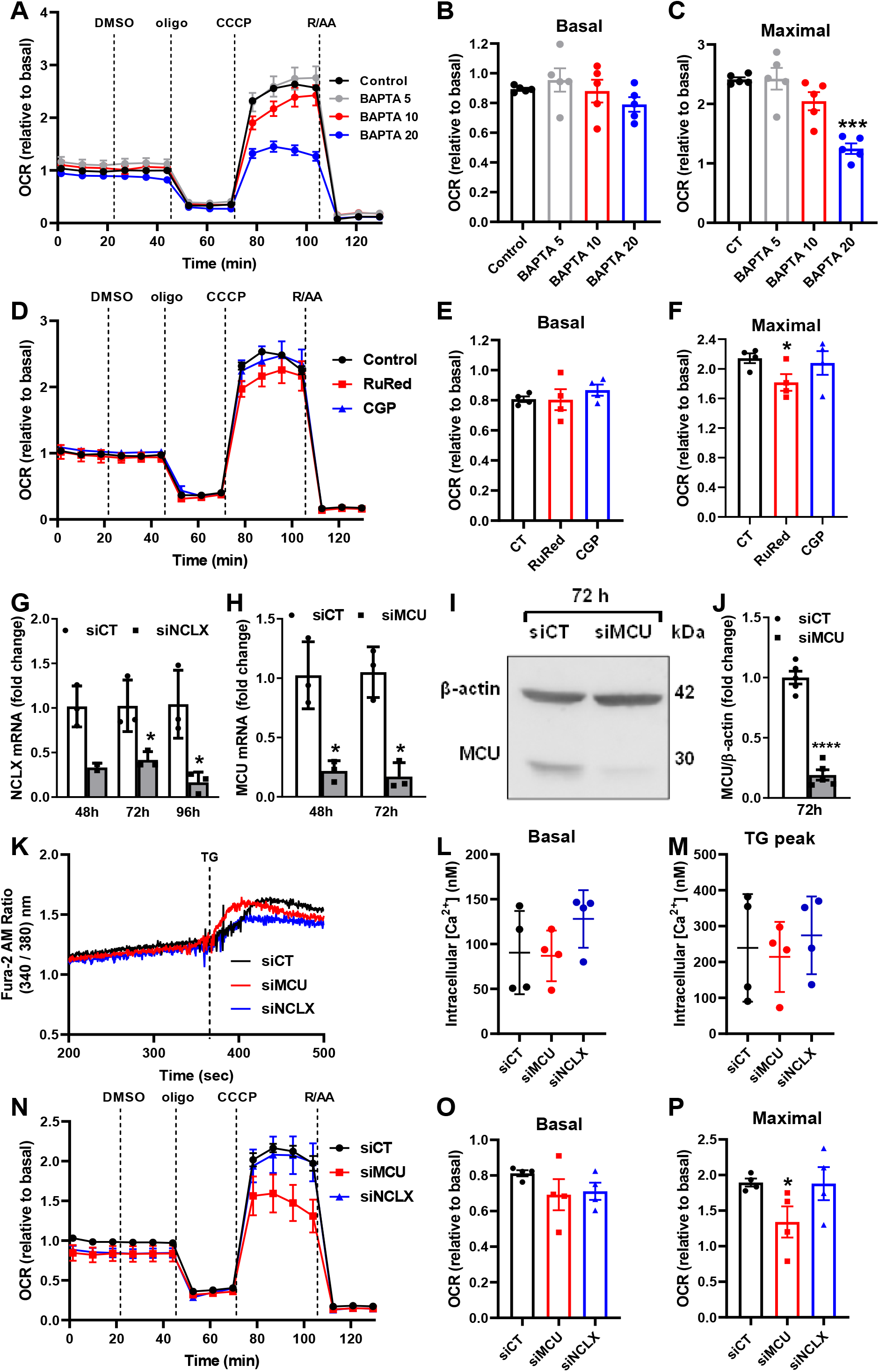
Depletion of intracellular Ca^2+^ stores and inhibition of mitochondrial Ca^2+^ uptake in intact cells limits maximal mitochondrial respiration. **A-C)** AML12 cells were pre-incubated for 1 h in the absence (control) or presence of an intracellular Ca^2+^ chelator, BAPTA-AM, at different concentrations (5 - 20 μM, as shown), and OCRs were measured using a Seahorse Extracellular Flux Analyzer. **D-F)** AML12 cells were incubated for 1 h in the absence (control, DMSO) or presence of 10 μM of RuRed or CGP, and OCRs was measured using a Seahorse Extracellular Flux Analyzer. **G-J)** MCU and NCLX silencing. **G)** NCLX mRNA levels after 48, 72 and 96 h of transfection with specific siRNA against NCLX (siNCLX) or a negative control (siCT). **H)** MCU mRNA levels after 48 and 72 h of transfection with specific siRNA against MCU (siMCU) or a negative control (siCT). **I)** Representative western blots of MCU and β-actin (internal control) after 72 h of transfection with specific siRNA against MCU (siMCU) or a negative control (siCT). **J)** Densitometric analysis of the Western blots. K-M) Cytosolic Ca^2+^ levels in siMCU or siNCLX cells. **K)** Typical cytosolic Ca^2+^ measurements performed using Fura-2AM. **L)** Quantification under basal conditions. **M)** Quantification of the peak after the addition of thapsigargin (TG). **N-P)** OCRs of siCT, siMCU or siNCLX AML12 cells were measured using a Seahorse Extracellular Flux Analyzer. Oligomycin (oligo, 1 μM), CCCP (5 μM), and rotenone + antimycin (R/AA, 1 μM each) were added where indicated. **B,E,O)** Quantification of basal respiration. **C,F,P)** Quantification of maximal respiration. Results are expressed as means ± SEM of 3-4 independent experiments; *p<0.05 versus control using Student’s t-test. Results are expressed as representative traces **(A, D, K, N)** or means ± SEM of 3-5 independent biological experiments; *p<0.05, ***p<0.001 and ****p<0.0001 versus control using one-way ANOVA followed by Šidak or Student’s t-test.

To measure the respiratory effects of physiological Ca^2+^ concentrations, we removed increasing amounts of this ion from the cytosol by incubating AML12 cells in complete medium with an intracellular Ca^2+^ chelator, BAPTA-AM, at different concentrations (5, 10 and 20 μM) (Fig. 5A-C). BAPTA-AM reduced maximal OCRs (Fig. 5C), but no changes were observed in basal respiration (Fig. 5B), which in intact cells corresponds to the respiration necessary to maintain normal levels of oxidative phosphorylation for typical cell function, in addition to the proton leak and non-mitochondrial oxygen consumption (inhibited in each trace by the addition of rotenone plus antimycin A, R/AA). Thus, decreasing physiological intracellular Ca^2+^ levels hamper maximal electron transfer capacity in intact cells, but not sufficiently to decrease ATP-linked respiration under normal (basal) conditions.

We then questioned if the effects of physiological levels of intracellular Ca^2+^ on OCRs in intact cells required Ca^2+^ entry into the matrix, as we found in isolated mitochondria. Inhibition of the MCU or NCLX was promoted in AML12 cells both pharmacologically with RuRed or CGP, and by silencing with specific siRNAs. We found that an acute inhibition of the transporters (1 h in the presence of pharmacological inhibitors) did not influence basal respiration (Fig. 5D,E), but MCU inhibition with RuRed led to significantly decreased maximal respiration (Fig. 5D,F). Cells in which NCLX was inhibited by CGP had similar profiles to untreated control cells (Fig. 5D-F).

We then checked how prolonged inhibition of those channels impacted respiration by silencing MCU or NCLX with siRNAs (Fig. 5G-J). Interestingly, we found that overall cytosolic Ca^2+^ levels were similar in these cells, despite the lack of these major mitochondrial Ca^2+^ regulatory pathways, both under basal conditions and after ER Ca^2+^ depletion due to thapsigargin (TG) addition (Fig. 5K-M). Chronic inhibition of both channels did not significantly alter basal respiration (Fig. 5N,O), while MCU knockdown led to a significant decrease in maximal respiration (Fig. 5N,P). This shows that both short and long-term MCU inhibition impact on mitochondrial activity in intact hepatocyte-derived cells.

In order to increase physiological intracellular Ca^2+^ levels and study the effects on mitochondrial oxidative phosphorylation, we used two approaches: a physiological stimulus with adrenaline (ADR) and a sarco-/endoplasmic reticulum Ca^2+^ ATPase (SERCA) pump inhibitor, TG. The effects of ADR and other Ca^2+^-mobilizing hormones on mitochondrial metabolic responses have been previously investigated in the liver (26,32): ADR binds to Gαs receptor and activates adenylyl cyclase, increasing cyclic AMP (cAMP) and promoting IP3 generation, leading to Ca^2+^ release from intracellular stores such as the endoplasmic reticulum (ER) stores (33). As the ER and mitochondria are closely connected in microdomains called mitochondria-associated ER membranes, this is expected to impact on mitochondrial Ca^2+^ homeostasis.

We first checked for intramitochondrial Ca^2+^ levels after addition of TG or ADR (Fig. 6A). Although both strategies led to increased intramitochondrial Ca^2+^ levels, the kinetics of this effect were very different. TG induced a much higher increase in intramitochondrial Ca^2+^ than ADR. In addition, ADR induced a transient and moderate increase in intramitochondrial Ca^2+^, while intramitochondrial Ca^2+^ levels remained elevated and did not return to basal levels after TG (Fig. 6A).

**Figure 6.**
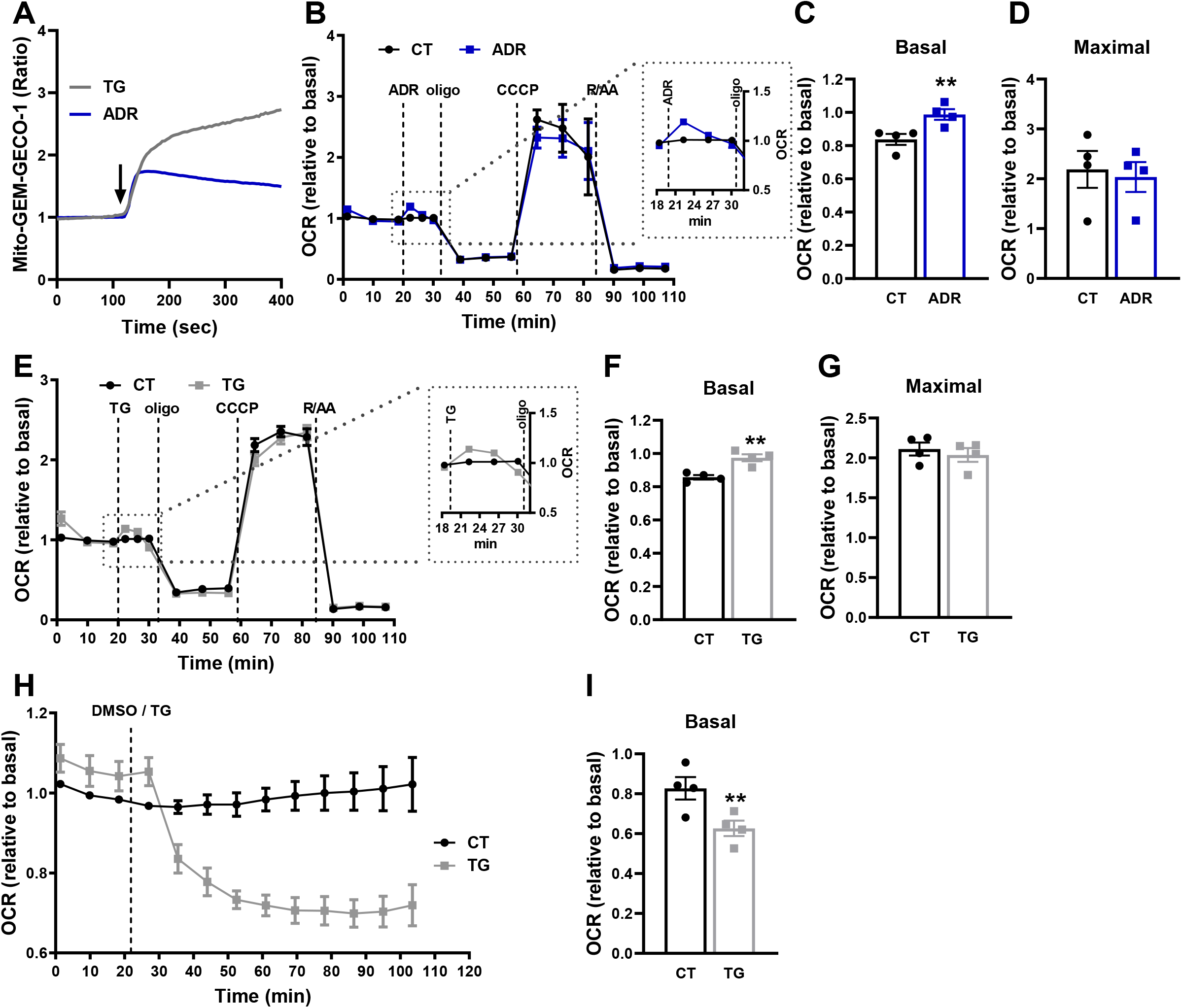
Dual effects of Ca^2+^ on mitochondrial respiration in intact cells. **A)** Representative images of mito-GEM-GECO1 AML12 cells. AML12 cells were transfected with mito-GEM-GECO1 for the evaluation of intramitochondrial Ca^2+^ levels. Fluorescence was assessed by confocal microscopy under basal conditions, followed by addition of thapsigargin (TG, 2 μM) or adrenaline (ADR, 20 μM). The arrow indicates ADR/TG addition. **B-I)** OCRs of AML12 cells were measured using a Seahorse Extracellular Flux Analyzer under basal conditions, followed by injection of DMSO (diluent), 20 μM ADR or 2 μM TG, oligomycin (oligo, 1 μM), CCCP (5 μM), and rotenone + antimycin (R/AA, 1 μM each). **B,E)** OCR traces. The insets represent the magnified area indicated by the dotted lines. **C,F)** Quantification of basal respiration. **D,G)** Quantification of maximal respiration. **H)** Long traces of OCRs after TG injection. **I)** Quantification of basal respiration of I. Results are expressed as means ± SEM of 4 independent experiments. **p<0.01 versus respective untreated control using Student’s t-test.

Interestingly, AML12 cells displayed an early increase in OCRs immediately after ADR injection (Fig. 6B), which was transient and rapidly went back to control levels. This ADR effect resulted in a significant increase in basal respiration (Fig. 6C), with no changes in maximal respiration (Fig. 6D). We observed a similar effect with TG: an early increase in OCRs a few minutes after TG injection (Fig. 6E), reflecting a significant increase in basal respiration (Fig. 6F), but with no changes in maximal respiration (Fig. 6G). As with ADR, the effect of TG was also transient, but instead of returning to basal conditions, OCR with TG started to decrease between 4-8 minutes after its injection (Fig. 6E). We then designed a long-term evaluation of OCRs after TG injection and found that the decreased OCR is permanently maintained and not reversed over time (Fig. 6H-I). Taken together, the results in Figs. 5–6 show that physiological intracellular Ca^2+^ homeostasis is maintained close to the Goldilocks “sweet spot” for oxidative phosphorylation activity, with both sustained increases and decreases in this concentration hampering ideal electron transport function.

### 2.3. Ca^2+^-induced impairment of mitochondrial respiration in situ is substrate-dependent

We next speculated if the Ca^2+^ effects on respiration in intact cells could also be substrate-dependent. We used three inhibitors that affect the oxidation of primary substrates that fuel mitochondria: UK5099, an inhibitor of the mitochondrial pyruvate carrier, MPC, which inhibits mitochondrial glucose/pyruvate oxidation; BPTES, an inhibitor of glutaminase 1, GLS-1; and etomoxir, an inhibitor of carnitine palmitoyl transferase 1a, CPT1a. We found that the decrease in basal and maximal respiration after thapsigargin was prevented with glucose/pyruvate metabolism inhibition by UK5099 (Fig. 7A,B), showing that the modulation of respiration by Ca^2+^ in intact cells is dependent on pyruvate oxidation, corroborating data obtained with isolated mitochondria (Fig. 1).

**Figure 7.**
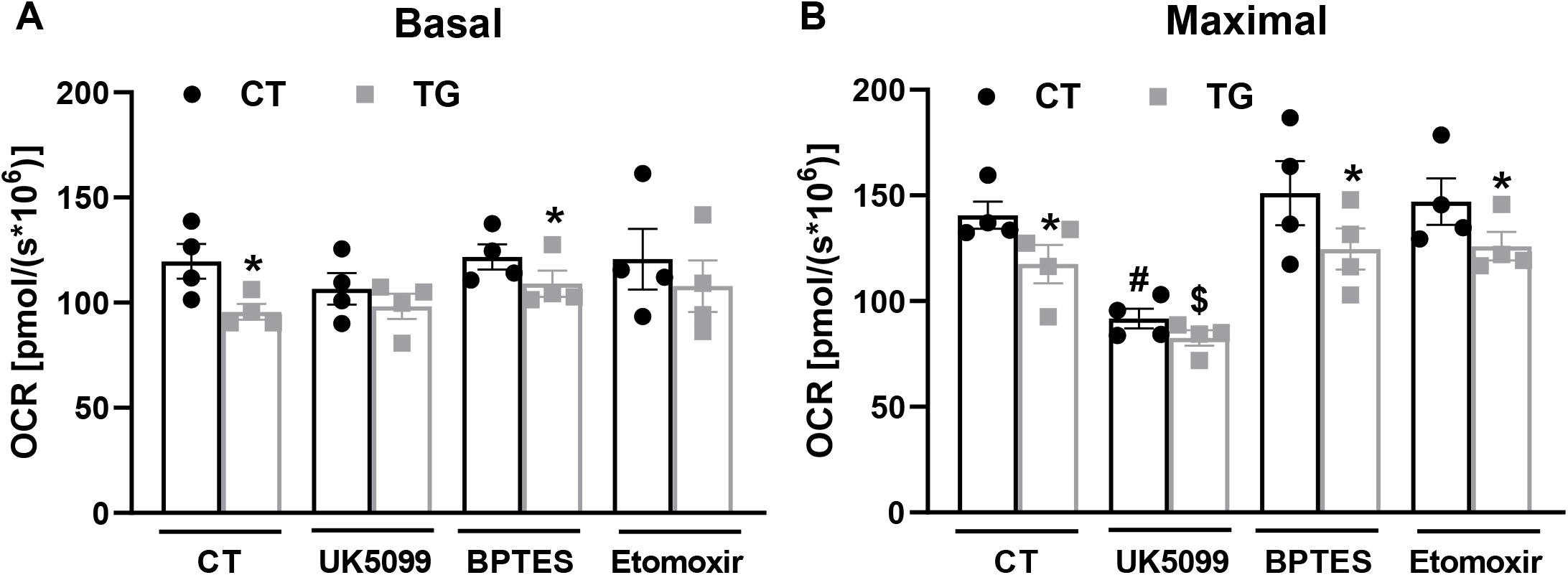
Intracellular Ca^2+^ modulates mitochondrial pyruvate oxidation. AML12 cells were incubated in the absence (CT) or presence of 2 μM thapsigargin (TG) and inhibitors (12 μM UK5099, 8 μM BPTES, or 14 μM Etomoxir). OCRs were measured using a high-resolution Oxygraph-2k (O2k) respirometer under basal conditions, followed by injection of oligomycin (oligo, 1 μM) and CCCP (5 μM), to induce maximal respiration. **A)** Quantification of basal respiration. **B)** Quantification of maximal respiration. Results are expressed as means ± SEM of 4 independent experiments. *p<0.05 versus respective untreated control using Student’s t-test. #p<0.05 versus CT, BPTES and Etomoxir under control conditions and $p<0.05 versus CT, BPTES and Etomoxir in TG condition, using one-way ANOVA followed by Šidak.

### 2.4. Ca^2+^-induced impairment of mitochondrial respiration in hepatocyte-derived cells occurs secondarily to mPTP opening

Having uncovered the effects of mitochondrial Ca^2+^ cycling on cellular respiration, we sought to dissect the reasons for prolonged decreased basal respiration induced by Ca^2+^ release from the ER promoted by TG (Fig. 8). Acute MCU inhibition with RuRed (Fig. 8A-D), as well as chronic MCU inhibition with MCU knockdown (Fig. 8E-H) protects mitochondrial respiration against TG-induced decreases, indicating that Ca^2+^ from the ER released by TG is taken up by mitochondria and affects electron transport.

**Figure 8.**
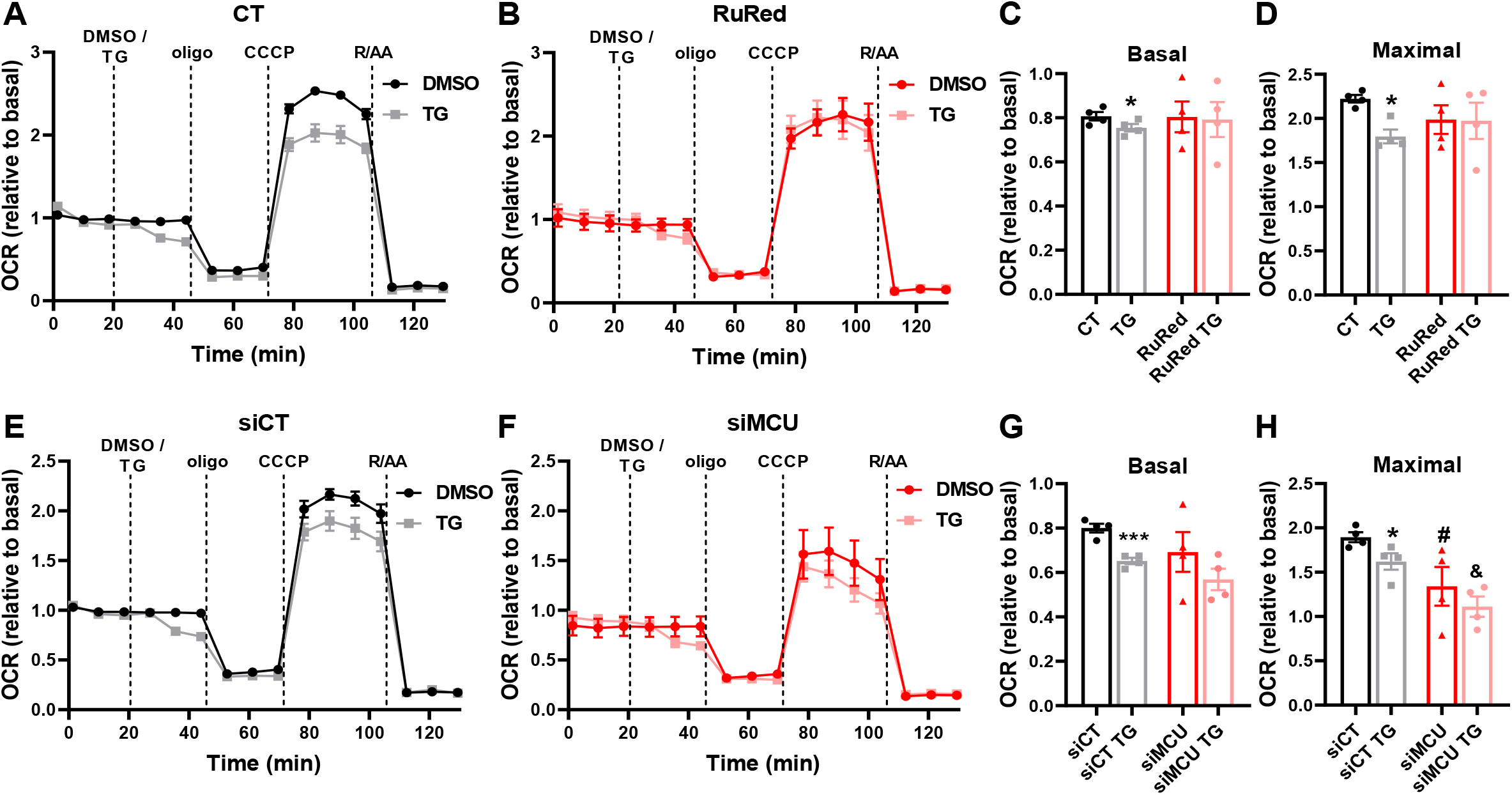
High intracellular Ca^2+^ limits maximal respiration in a manner dependent on mitochondrial Ca^2+^ uptake. **A-D)** AML12 cells were preincubated for 1 h in the absence (CT) or presence of 10 μM of RuRed or CGP. **E-H)** Cells treated with specific siRNAs against the MCU (siMCU) or NCLX (siNCLX) or a negative control (siCT). OCRs were measured in a Seahorse Extracellular Flux Analyzer under basal conditions, followed by injection of DMSO (diluent) or 2 μM thapsigargin (TG), oligomycin (oligo, 1 μM), CCCP (5 μM), and rotenone + antimycin (R/AA, 1 μM each).**C,G)** Quantifications of basal and **(D,H)** maximal respiration. Results are expressed as mean ± SEM of 4 independent experiments; *p<0.05 and ***p<0.001 versus respective control using Student’s t-test, #p<0.05 versus siCT, & p<0.05 versus siCT TG using one-way ANOVA followed by Šidak.

Next, we analyzed intramitochondrial Ca^2+^ levels in AML12 cells transfected with a specific mitochondrial Ca^2+^ sensor (mito-GEM-GECO1) (34). This sensor has a Ca^2+^-dependent increase in fluorescence, and intramitochondrial Ca^2+^ changes can be easily visualized by changes in color. We found a rapid increase in intramitochondrial Ca^2+^ after TG addition (Fig. 9A-C), which was prevented by preincubation with the intracellular Ca^2+^ chelator BAPTA-AM (Fig. 9-C). In line with this, BAPTA-AM also protected against the TG-induced decrease in mitochondrial respiration (Fig. 9E,H,I).

**Figure 9.**
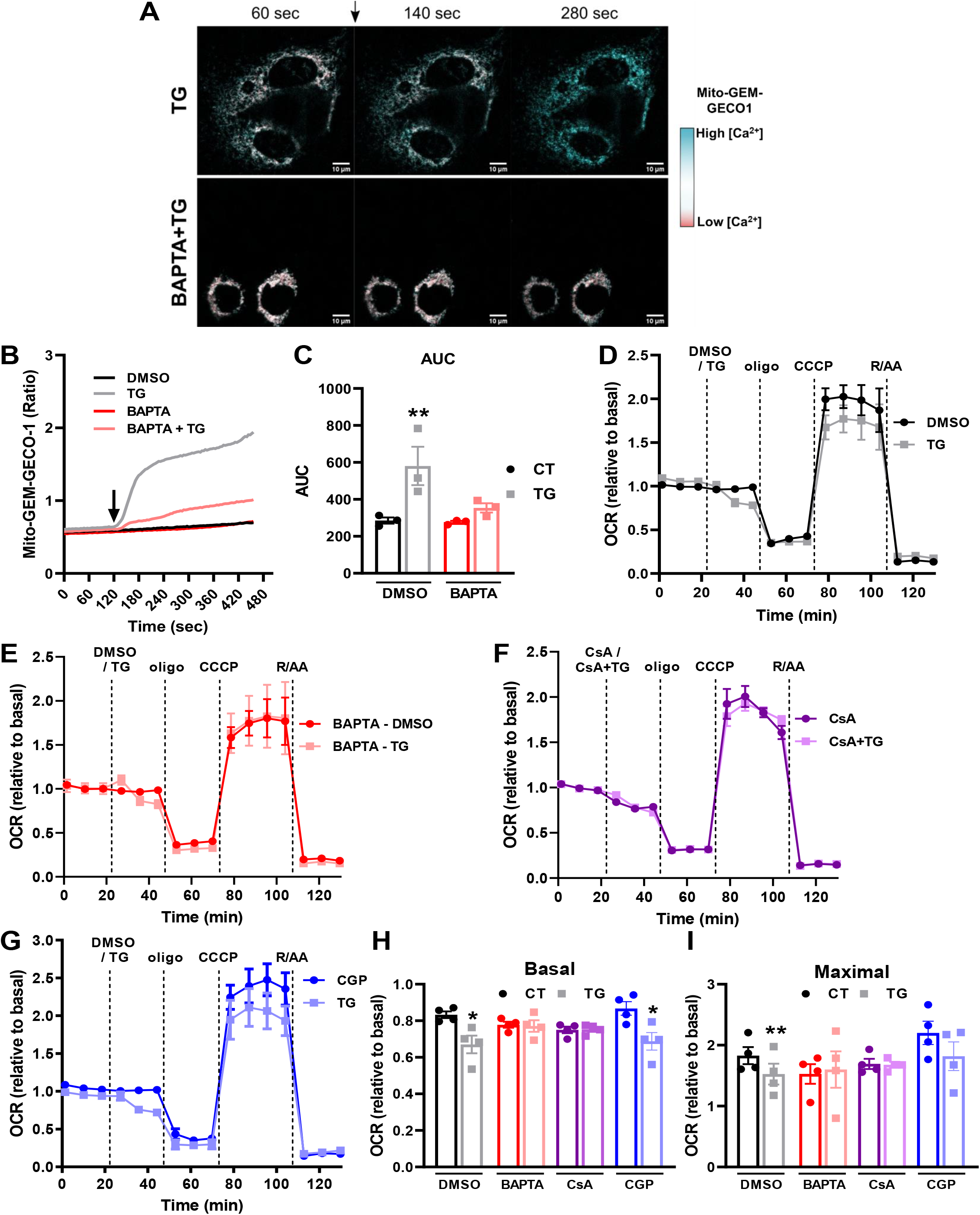
Respiratory inhibition promoted by excess Ca^2+^ is due to mPTP. **A-C)** AML12 cells were transfected with mito-GEM-GECO1 for the evaluation of intramitochondrial Ca^2+^ levels. Cells were preincubated in the absence or presence of 10 μM BAPTA-AM, and fluorescence was assessed by confocal microscopy under basal conditions, followed by addition of diluent (DMSO) or thapsigargin (TG, 2 μM). **A)** Representative images of mito-GEM-GECO1 AML12 cells. The arrow indicates TG addition. Cyan indicates high intramitochondrial [Ca^2+^], while red indicates low intramitochondrial [Ca^2+^]. **B)** Representative trace. **C)** Area under the curve (AUC) of B. **D-G)** OCRs were measured using a Seahorse Extracellular Flux Analyzer under basal conditions followed by injection of i) DMSO (diluent), 2 μM thapsigargin (TG), 10 μM cyclosporin A (CsA) or TG + CsA, ii) oligomycin (oligo, 1 μM), iii) CCCP (5 μM), and iv) rotenone + antimycin (R/AA, 1 μM each). **E)** Cells were pre-incubated for 1 h in the absence or presence of an intracellular Ca^2+^ chelator, 10 μM BAPTA-AM (BAPTA), before OCR measurements. **G)** Cells were pre-incubated for 1 h in the absence or presence of 10 μM CGP-37157 (CGP), before OCR measurements. **H)** Quantification of basal respiration. **I)** Quantification of maximal respiration. Results are expressed as mean ± SEM of 4 independent experiments; *p<0.05 and **p<0.01 versus respective control using Student’s t-test.

Ca^2+^ overload in mitochondria is known to promote opening of the mPTP (18–20), which can decrease OCRs by limiting mitochondrial cytochrome c content and affecting substrate import into the organelle. Thus, we incubated cells with TG in the presence of the mPTP inhibitor CsA, which efficiently prevented TG effects (Fig. 9F,H,I), showing the involvement of mPTP in TG-induced OCR decrease. We then blocked mitochondrial Ca^2+^ efflux through NCLX with CGP, and found it did not prevent TG-induced OCR decrease (Fig. 9G-I), reinforcing the role of mPTP and corroborating the results found in isolated mitochondria showing that CGP induces mPTP formation, leading to a decrease of calcium retention capacity and negatively impacting mitochondrial respiration.

## 3. Discussion

Mitochondria are well recognized as a major hub for calcium (Ca^2+^) handling and energy metabolism (1). Small amounts of Ca^2+^ have been shown to increase the affinity of some matrix dehydrogenases for their substrates, and thus assumed to possibly positively impact on ATP production. This assumption is a tricky one, since Ca^2+^ acts in different ways in mitochondria depending on concentration. In this sense, it has been previously shown that primary hepatocytes exposed to hormones such as vasopressin have increased cytosolic Ca^2+^ levels, which induce a subsequent increase in mitochondrial Ca^2+^ levels and enhanced activity of mitochondrial dehydrogenases (26,27). However, interestingly, both slow/partial and sustained increases in cytosolic Ca^2+^ levels were ineffective in modulating these processes (27). In addition, others have shown that conditions that induce excessive mitochondrial Ca^2+^ accumulation, such as the combination of vasopressin and glucagon, or thapsigargin, can be deleterious to hepatocytes (25). Indeed, matrix Ca^2+^ overload has been shown to induce mitochondrial swelling (24,25), and to promote impairment of mitochondrial function and cell death (35) in a myriad of pathological conditions.

We designed two types of experiments aimed at understanding the effects of Ca^2+^ ions on oxidative phosphorylation in a more global manner, both focused on liver metabolism, as the effects of different calibrated amounts of Ca^2+^ ions on this organ’s rich metabolic activities have not been extensively studied with respect to mitochondrial electron transport. We mainly sought to compare the effect of low versus supraphysiological increases in cytosolic and mitochondrial Ca^2+^ on oxygen consumption rates, using careful quantitative studies and modern methodology. The first approach involved isolated mitochondria (Figs. 1–4), in which precise concentrations of extramitochondrial Ca^2+^ can be added, and the metabolism of different substrates can be examined while still preserving intact oxidative phosphorylation pathways, thus allowing to establish the impact of Ca^2+^ ions on overall metabolic fluxes.

Our results show that isolated liver mitochondria present enhanced oxidative phosphorylation efficiency promoted by Ca^2+^, in a manner dependent on the substrate used and Ca^2+^ concentrations. Increased respiratory control ratios (RCRs) were achieved with low extramitochondrial Ca^2+^ (2.4 ± 0.6 μM) additions in the presence of the complex I substrates pyruvate plus malate (Fig. 1), and with medium Ca^2+^ (22.0 ± 2.4 μM) in the presence of α-ketoglutarate (Fig. 2). Interestingly, previous reports show that pyruvate dehydrogenase (PDH) and α-ketoglutarate dehydrogenase (α-KGDH) are directly or indirectly activated by Ca^2+^ within the same concentration range (0.1 to 10 μM) in rat mitochondria from heart (30,31), skeletal muscle (32), adipose tissue (33), and liver (34). However, our results suggest that more Ca^2+^ is required to change α-KGDH-versus PDH-supported respiration in mouse liver mitochondria (Figs. 1,2). This confirms that results showing changes in affinities of isolated enzymes cannot be used as predictors of metabolic fluxes in intact organelles. Indeed, changes in affinity cannot predict metabolic fluxes without considering the concentrations of the substrates/intermediates, as enzymes activated by Ca^2+^ may already be at their Vmax in the microenvironment of the mitochondrial matrix.

On the other hand, when complex II substrate succinate was used, extramitochondrial Ca^2+^ did not modulate mitochondrial efficiency (Fig. 3). In line with this, recent results (36) show that isolated heart and kidney mitochondria exposed to pyruvate plus malate or α-ketoglutarate plus malate have increased RCRs when in the presence of small increases in Ca^2+^ concentrations (in the nanomolar range), but had decreased RCRs when Ca^2+^ was further augmented. In contrast, Ca^2+^ addition to isolated brain mitochondria energized by pyruvate plus malate leads to reduced RCRs, in a concentration-dependent manner, with no effect on PDH activity (37). These results, overall, show that positive effects of Ca^2+^ on respiration are highly tissue-dependent, and, when present, occur within a tight range of concentrations.

With the exception of some careful studies showing intracellular Ca^2+^ quantifications followed by hormone-induced mitochondrial Ca^2+^ increases (23), a major concern regarding mitochondrial Ca^2+^ effects presented in the literature is a possible lack of homogeneity in Ca^2+^ concentrations used not only between different studies, but also in replicates within each study. Different Ca^2+^ amounts can be present in distinct isolated mitochondrial preparations, as these carry with them varying quantities of chelators from the isolation buffer. In order to minimize possible confounding results, we found it crucial to determine the amount of Ca^2+^ present after isolation daily, and adjust Ca^2+^ added, so it was equal between biological replicates. We did this using Ca^2+^ uptake assays, as described in the methods section. This daily calibration may have been a seminal feature which allowed us to uncover activating Ca^2+^ effects, as these were often subtle and concentration-specific. We also exclusively tested Ca^2+^ concentrations in which we observed no mitochondrial permeability transition pore (mPTP) opening, as indicated by the fact that mitochondrial preparations were able to uptake and retain the ion. Interestingly, the concentrations necessary to induce mPTP are different with distinct substrates, and larger for succinate (Fig. 3A) than NADH-linked substrates (Figs. 1A and 2A), thus allowing us to test more Ca^2+^ concentrations on respiration supported by succinate (with negative results).

While isolated mitochondrial studies allow the dissection of precise effects of substrates and specific ion concentrations, they do not represent the metabolic state of oxidative phosphorylation in intact cells. Thus, our second approach was to modulate mitochondrial Ca^2+^ levels in the non-tumor AML12 hepatocyte-derived cell line (30). In intact cells, overall cytosolic Ca^2+^ concentrations are in the 100-150 nM range (Fig. 5K-L), which is much lower than concentrations that activate oxidative phosphorylation in isolated mitochondria, but does not reflect differences in local ion concentrations, such as those in the Ca^2+^-rich mitochondrial-ER contact sites, where Ca^2+^ concentrations may be up to 10 times higher than in the bulk cytosol (38–40).

We found that the Ca^2+^ concentrations in mitochondria *in situ* are both necessary and sufficient to enhance electron transport capacity through various different approaches. First, chelation of intracellular Ca^2+^ with BAPTA-AM decreased maximal OCRs (Fig. 5C), without affecting mitochondrial membrane integrity, as indicated by the lack of change in oligomycin-insensitive respiration (Fig. 5A). Second, inhibition of mitochondrial Ca^2+^ uptake with ruthenium red (RuRed; Fig. 5D,F) or by silencing the mitochondrial Ca^2+^ uniporter (MCU) (Fig. 5N,P) also decreased maximal electron transport capacity in intact cells, indicating that ideal respiratory chain function requires both the presence of physiological levels of cytosolic Ca^2+^ and its uptake by mitochondria. Our results are compatible with data showing that genetic manipulation of MCU and consequent impairment of mitochondrial Ca^2+^ uptake disrupts oxidative phosphorylation and lowers cellular ATP in HeLa cells (39). Similarly, mitochondria from a liver-specific MCU knockout mouse model also had inhibited mitochondrial Ca^2+^ uptake and reduced oxidative phosphorylation (40).

Conversely, we induced ER Ca^2+^ depletion with adrenaline (ADR), promoting IP3-mediated ER Ca^2+^ release, and by adding the sarco-/endoplasmic reticulum Ca^2+^ ATPase (SERCA) pump inhibitor thapsigargin (TG). Both IP3R and SERCA are located in mitochondrial-associated ER membrane microdomains where the organelles are in close proximity, allowing for Ca^2+^ exchange (41,42). We show that increased cytosolic Ca^2+^ in intact cells (with TG or ADR) induces a moderate (with ADR) or sustained (with TG) increase in intramitochondrial Ca^2+^ levels, leading to an early and transient stimulation of respiratory rates, followed by a decrease to control levels (with ADR) and an even further decrease with TG (Fig. 6), showcasing the effects over time and of different mechanisms and intensities of Ca^2+^ increases. Altogether, our results indicate that physiological Ca^2+^ levels in mitochondria are close to the ideal level to maximize electron transport, with both sustained decreases and increases leading to lower efficiency in this process. These results are in line with the finding that constitutive IP3R-mediated ER Ca^2+^ release to mitochondria is essential for efficient mitochondrial respiration and maintenance of normal cell bioenergetics (43). Ca^2+^ release from the ER, both through ryanodine receptors (RYRs) and IP3Rs, has also been shown to elevate matrix Ca^2+^ levels and stimulate mitochondrial oxidative phosphorylation in neurons (44). On the other hand, cells expressing a truncated variant of SERCA-1 with increased ER-mitochondria contact sites and increased Ca^2+^ transfer from the ER to mitochondria show signs of mitochondrial dysfunction due to Ca^2+^ overload (45).

Maximal OCR limitation observed under these conditions of Ca^2+^ overload in intact cells is due to uptake of the ion into the mitochondrial matrix, as it is inhibited by BAPTA-AM, MCU silencing or pharmacological inhibition (Figs. 8 and 9). Excess mitochondrial Ca^2+^ is known to lead to mPTP opening and consequent dissipation of the mitochondrial membrane potential, mitochondrial swelling and rupture, culminating in cell death (18–20). Since the decrease in OCR observed under our conditions of mitochondrial Ca^2+^ overload was prevented by mPTP inhibitor cyclosporin A (CsA, Fig. 9), it is attributable to this form of mitochondrial inner membrane permeabilization. As we did not detect irreversible mitochondrial injury after TG-induced mPTP, we speculate that this may be due to transient or perhaps flickering mPTP, a consequence of mitochondrial Ca^2+^ loading above normal physiological levels.

While it may seem instinctively obvious that physiological resting mitochondrial Ca^2+^ levels would be at ideal quantities to activate maximum electron transport, but not generate mitochondrial damage, it should be noted that not all intracellular signaling situations operate physiologically at maximum capacity. Perhaps the best example in this context is mitochondrial electron transfer itself, which is typically lower under basal conditions than maximum capacity uncovered when in the presence of uncoupler (see Figs. 5–9). Indeed, the difference between basal OCR and maximal achievable electron transport, known as reserve capacity, is thought to be important in resistance toward stress; a manner to allow fast increments in ATP synthesis when necessary (46). On the other hand, our results show clearly that physiological Ca^2+^ levels in mitochondria are specifically tailored to maintain the highest maximal OCR without promoting mPTP. Overall, we demonstrate that hepatocyte-derived cells maintain cellular and mitochondrial Ca^2+^ concentrations within strictly controlled ranges, maximizing electron transport capacity by promoting the Goldilocks Ca^2+^ “sweet spot” for oxidative phosphorylation activity: neither too much, nor too little, but just right.

## 4. Experimental procedures

### 4.1. Reagents

Culture medium (DMEM/F-12) (#11320), fetal bovine serum (FBS), insulin-transferrin-selenium (ITS), Ca^2+^ Green 5N and Pierce BCA protein assay kit were from Thermo Fischer Scientific (Waltham, MA, USA); thapsigargin, adrenaline, safranin-O, BAPTA-AM, CGP-37157, ruthenium red, UK5099, BPTES, etomoxir and RIPA buffer were from Sigma-Aldrich (St. Louis, MO, USA); lipofectamine RNAiMAX, lipofectamine 2000, Trizol reagent and Fura-2 AM were from Invitrogen (Waltham, MA, USA); CMV-mito-GEM-GECO1 plasmid (#32461) was from Addgene (Watertown, MA, USA); Bradford was from Bio-Rad Laboratories (Hercules, CA, USA); siRNAs against MCU (ID s103465), NCLX (ID s100747) or negative control (#4390844) were from Ambion Inc. (Austin, TX, USA); anti-MCU (#14997S) was from Cell Signaling (Danvers, MA, USA); anti-β-actin (#ab8226) was from Abcam (Cambridge, UK); fluorescent secondary antibodies (goat anti-mouse #926-68070 and goat anti-rabbit #926-68071) were from Licor (Lincoln, NE, USA).

### 4.2. Animals

C57BL/6NTac male mice were bred and housed in the animal facility of the Faculty of Pharmaceutical Sciences and Chemistry Institute of the University of São Paulo, devoid of murine pathogens. Animals were maintained in collective cages (max 4/cage) at controlled temperatures (23°C) in a 12–12 h light/dark cycle with free access to food/water. We used 10–12-week-old mice. All procedures were conducted in accordance with the Ethical Principles of Animal Manipulation of the local animal ethics committee, under protocol CEUA-IQ/USP 196/2021.

### 4.3. Isolation of liver mitochondria

After deep anesthesia (4% isoflurane) followed by cervical dislocation, the abdomen was dissected and the liver was removed. Mitochondria were isolated by differential centrifugation. All steps were carried out at 4°C, over ice. Briefly, the liver was chopped into small pieces, suspended in isolation buffer (250 mM sucrose, 10 mM HEPES, 1 mM EGTA, 1 mM EDTA, 1 mg/mL BSA, pH 7.2), and manually homogenized using a Potter-Elvehjem tissue grinder. To remove blood from the mixture, it was centrifuged twice (800 g, 4°C, 4 min), and the pellet was discharged. The supernatant was further centrifuged at 9,000 g, 4°C for 10 min. The pellet was resuspended in resuspension buffer (300 mM sucrose, 10 mM HEPES, and 2 mM EGTA, pH 7.2), and centrifuged again (9,000 g, 4°C, 10 min). The pellet was resuspended in 125 μL of resuspension buffer, and total protein was quantified using the Bradford method.

### 4.4. Oxygen consumption by isolated liver mitochondria

Oxygen consumption was measured using a high-resolution Oxygraph-2k (O2k) respirometer (Oroboros Instruments, Innsbruck, Austria). Mitochondria (600 μg protein for pyruvate/malate and α-ketoglutarate; 200 μg protein for succinate/rotenone) were incubated in 2 mL of experimental buffer (125 mM sucrose, 65 mM KCl, 10 mM HEPES, 2 mM MgCl_2_, 2 mM KH_2_PO_4_, and 0.1 mg/mL BSA) containing 100 μM EGTA and different substrates (1 mM pyruvate + 1 mM malate, 1 mM α-ketoglutarate or 1 mM succinate + 1 μM rotenone) and/or inhibitors (10 μM ruthenium red or 10 μM CGP-37157) at 37°C with continuous stirring (700 rpm). After basal respiration was measured, CaCl_2_ was injected at different concentrations, as indicated. After reaching a steady state (as indicated by Ca^2+^ uptake measurements), state 3 respiration was measured by adding 1 mM ADP, followed by addition of 1 μM oligomycin (State 4), and 0.5 μM CCCP (state 3u). Respiratory control ratios (RCR) were calculated as State 3/State 4.

### 4.5. Calcium (Ca^2+^) uptake assays

Ca^2+^ measurement assays were performed simultaneously with oxygen consumption measurements, in order to determine the approximate free extramitochondrial Ca^2+^ concentration and to monitor Ca^2+^ levels over time, as done in (47), as well as to monitor mPTP opening. Mitochondria (600 μg protein for pyruvate/malate and α-ketoglutarate; 200 μg protein for succinate/rotenone) were incubated in 2 mL of the same experimental buffer used in the oxygen consumption assay, containing 0.1 μM Calcium Green 5N, 100 μM EGTA and different substrates and inhibitors, as indicated. Calcium Green fluorescence was measured at λ_ex_ = 506 nm and λ_em_ = 532 nm, using a F4500 and a F2500 Hitachi Fluorimeters at 37°C with continuous stirring. After a 100 s baseline interval, a CaCl_2_ solution was added to achieve the target Ca^2+^ concentrations: low Ca^2+^ (added Ca^2+^ at 30-50 μM or 0.1-0.17 nmol Ca^2+^/μg mitochondria for pyruvate/malate and α-ketoglutarate and 0.3-0.5 for succinate/rotenone), medium Ca^2+^ (added Ca^2+^ at 45-75 μM or 0.15-0.25 nmol Ca^2+^/μg mitochondria for pyruvate/malate and α-ketoglutarate and 0.45-0.75 for succinate/rotenone) and high Ca^2+^ (added Ca^2+^ at 75-100 μM or 0.25-0.33 nmol Ca^2+^/μg mitochondria for pyruvate/malate and α-ketoglutarate and/or 0.75-1.0 for succinate/rotenone). According to the amount of Ca^2+^ added and the excess EGTA present in the buffer, we calculated the amount of free extramitochondrial Ca^2+^ to which mitochondria were exposed: low (2.4 ± 0.6 μM), medium (22.0 ± 2.4 μM) and high (52.9 ± 2.5 μM) concentrations, from now on called simply as low, medium or high Ca^2+^. Fluorescence was monitored for 20 min. Each trace was followed by three 10 μM Ca^2+^, three 1 mM Ca^2+^, and three 3 mM EGTA additions, to allow the estimation of the Calcium Green 5N experimental Kd, and maximal (Fmax) and minimal (Fmin) fluorescence, respectively. Absolute fluorescence values (F) were calibrated into [Ca^2+^] through the formula [Ca^2+^] = Kd.(F - Fmin)/(Fmax - F). To determine Ca^2+^ retention capacity (CRC), several additions of 10 μM Ca^2+^ were made until mPTP opening occurred. These traces were performed with and without 5 μM CsA, as indicated.

### 4.6. Mitochondrial membrane potential measurements

Mitochondrial membrane potentials (ΔΨm) were determined by measuring changes in the fluorescence of safranin-O, in quenching mode. Mitochondria (600 μg protein) were incubated in 2 mL of the same experimental buffer used in 4.4. and 4.5., containing 5 μM safranin-O, 100 μM EGTA, and 1 mM pyruvate + 1 mM malate or 1 mM α-ketoglutarate. Fluorescence was measured at λ_ex_ = 485 nm and λ_em_ = 586 nm, using a F4500 Hitachi Fluorimeter at 37°C with continuous stirring. After a 100 s baseline interval, CaCl2 was added to achieve a low or medium Ca^2+^ concentration, and fluorescence was monitored for 20 min. Fluorescence values were calibrated to mV as described in (48,49). Briefly, ΔΨm was clamped at different time points using valinomycin (a K^+^ ionophore) and sequential additions of known KCl concentrations, and fluorescence was followed. A calibration curve was constructed based on fluorescence variations and estimated ΔΨm, calculated from the Nernst equation, ΔΨm = 60 × log(K^+^_in_/K^+^_out_), considering the intramitochondrial [K^+^]_in_ = 120 mM and known extramitochondrial K^+^ concentrations (K_out_).

### 4.7. NAD(P)H measurements

NAD(P)H was determined by measuring changes in the NAD(P)H autofluorescence. Mitochondria (600 μg protein) were incubated in 2 mL of the same experimental buffer used in 4.4. and 4.5. and 4.6., containing 100 μM EGTA and different substrates, as indicated. Fluorescence was measured at λ_ex_ = 366 nm and λ_em_ = 450 nm, using a F2500 Hitachi Fluorimeter at 37°C with continuous stirring. After a 100 s baseline interval, a CaCl2 solution was added to achieve the target Ca^2+^ concentrations, as indicated, followed by sequential additions of 1 mM ADP, 1 μM oligomycin, 0.5 μM CCCP, and 1 μM rotenone + 1 μM antimycin A. Fluorescence values were calibrated to nM through a calibration curve constructed in experimental buffer, in absence of mitochondria, following additions of known NADH concentrations.

### 4.8. AML-12 cell cultures

The hepatocyte-derived AML-12 cell line was maintained in DMEM/F-12 medium, which contains 17.5 mM glucose, 2.5 mM glutamine, and several other amino acids, supplemented with 10% (v/v) FBS, a mixture of insulin, transferrin, and selenium (ITS; Collaborative Research), 1% antibiotics (100 U/mL penicillin, 0.1 mg/mL streptomycin), pH 7.4, at 37°C in a humidified atmosphere of 5% CO_2_.

### 4.9. Small interfering RNA (siRNA) transfection

AML-12 cells were transfected with siRNAs against MCU, NCLX or scramble RNA as a negative control (Ambion Inc.), using Lipofectamine RNAiMAX and a reverse transfection protocol. Briefly, lipofectamine and siRNAs were diluted in Opti-MEM medium and were added to the cell suspension at a final concentration of 20 nM. Cells were kept overnight at 37°C in DMEM/F-12 containing ITS, without antibiotics. After transfection, the medium was replaced by complete medium. 48 h after transfection, cells were seeded for experiments and left for an additional 24 h.

### 4.10. RT-qPCR

After transfections with siRNAs, AML-12 cells were collected with Trizol™ Reagent and RNA was isolated following the manufacturer’s instructions. Total RNA was quantified using a NanoDrop^®^ spectrophotometer and cDNA synthesis was performed using the High-Capacity cDNA Reverse Transcription Kit. qPCR reactions were performed using the Platinum^®^SYBR^®^ Green qPCR SuperMix-UDG with specific primers for *Mcu* (FW 5’-ACTCACCAGATGGCGTTCG-3’; RV 5’-CATGGCTTAGGAGGTCTCTCTT-3’), *Nclx* (FW 5’-TGTCACCTTCCTGGCCTTTG-3’; RV 5’-CACCCCTGCACCAAACAGA-3’), *Hmbs* (FW 5’-CAGCTACAGAGA AAGTTCCCC-3’; RV 5’-AGGACGATGGCACTGAATTC-3’) and *B2m* (FW 5’-CTGGTCTTTCTATATCCTGGCTC-3’; RV 5’-TGCTTGATCACATGTCTCGATC-3’) genes. Amplification data was analyzed by the 2^(-ΔΔCt)^ method using the mean of *Hmbs* and *B2m* Ct values as housekeeping. The housekeeping choice was based on the stability value of the gene or combination of genes calculated by the NormFinder Software (50).

### 4.11. Western blots

In order to validate MCU silencing, we performed a western blot analysis of MCU levels in transfected AML-12 cells. Briefly, cells were lysed in RIPA buffer containing protease and phosphatase inhibitor cocktails, and total protein levels were quantified using Pierce BCA Protein Assay kit. Lysates were prepared in sample buffer (2% SDS, 10% Glycerol, 0.062 M Tris pH 6.8, 0.002% Bromophenol Blue, 5% 2-mercaptoethanol), loaded onto SDS-PAGE gels and electrotransfered to PVDF membranes. Membranes were blocked with 5% BSA in TTBS (20 mM Tris pH 7.5, 150 mM NaCl and 0.1% Tween 20) for 1 hour at room temperature before overnight incubation with MCU (1:1,000) and β-actin (1:5,000) primary antibodies at 4°C. Fluorescent secondary antibodies (1:15,000) were incubated for 1 hour at room temperature prior to fluorescence detection using a ChemiDoc™ Imaging System (Bio-Rad). Quantification of band densitometry was performed using the FIJI ImageJ software (51).

### 4.12. Cytosolic Ca^2+^ measurements

After MCU or NCLX silencing, AML-12 cells were seeded at 1.5 × 10^6^ cells in 4 mL complete medium in 60 mm cell culture dishes. After 24 h, cells were loaded with 5 μM Fura-2 AM for 1 h at 37°C. After the loading period, cells were trypsinized and suspended in Krebs-Henseleit buffer without CaCl_2_ (115 mM NaCl, 24 mM NaHCO_3_, 5 mM KCl, 1 mM MgSO_4_, 1.2 mM KH_2_PO_4_). 1 × 10^6^ cells (final volume: 2 mL) were checked for basal cytosolic Ca^2+^ levels and after ER Ca^2+^ depletion with thapsigargin using a F4500 Hitachi Fluorimeter at 37°C with continuous stirring. Fluorescence was measured at excitation 340/380 nm and emission 505 nm. Each trace was followed by 4% Triton X-100 (50 μL) and 60 mM EGTA (150 μL) additions, to allow the estimation of maximal (R_max_) and minimal (R_min_) fluorescence ratio, respectively. Intracellular Ca^2+^ concentrations, [Ca^2+^]_i_, were calculated as described in (52) through the formula [Ca^2+^]_i_ (nM) = Kd × [(R –R_min_) / R_max_ – R)] × Sfb, where the K_d_ for Ca^2+^ binding to Fura-2 at 37°C = 225 nM and Sfb is the ratio of baseline fluorescence at 380 nm.

### 4.13. Intramitochondrial Ca^2+^ measurements

Intramitochondrial Ca^2+^ was measured in AML-12 cells expressing the mito-GEM-GECO1 sensor. Cells were seeded at 50,000 cells in 500 μL complete medium on 30 mm cell culture glass dishes, previously coated with poly-D-Lysine. After 24 h, cells were transfected overnight at 37°C in DMEM/F-12 without antibiotics containing Lipofectamine 2000 in the presence of a plasmid encoding a mitochondrial-targeted sequence of the Ca^2+^ sensor GEM-GECO1 (34). Cells were checked for fluorescence using a Confocal Zeiss LSM 780-NLO microscope at 37°C under different conditions: i) control media; ii) acute thapsigargin or adrenaline addition; iii) after 1h incubation with BAPTA-AM; iv) acute thapsigargin addition after 1h incubation with BAPTA-AM. Fluorescence was measured every 4 seconds with excitation at 390 nm and emission at 455/511 nm. Images were further analyzed using ImageJ FIJI Software (51). Results are presented as fluorescence of mito-GEM-GECO1 and the respective area under the curve (AUC).

### 4.14. Oxygen consumption in intact AML-12 cells

Cells were seeded at 30,000 cells per well in 100 μL DMEM/F-12 medium (with 10% FBS, 1% P/S, ITS) on Agilent Seahorse XF24 cell culture microplates and allowed to adhere overnight. Oxygen consumption was measured using a Seahorse Extracellular Flux Analyzer (Agilent Technologies, Santa Clara, CA, EUA). For that, cells were washed thrice with 500 μL DMEM/F-12 containing 1% P/S and 5 mM HEPES. Media did not contain bicarbonate nor FBS. Cells were kept for 1 h in a humidified incubator at 37°C without CO_2_, in the absence or presence of different additions (BAPTA-AM, 10 μM ruthenium red, 10 μM CGP-37157), as indicated. After incubation, plates were placed in the equipment and oxygen consumption rates (OCR) were measured under basal conditions, followed by different injections: a) thapsigargin (TG, final concentration: 2 μM) or same volume of DMSO in control wells or adrenaline (ADR, final concentration: 20 μM); b) oligomycin (final concentration: 1 μM); c) CCCP (final concentration: 5 μM); d) antimycin and rotenone (R/AA, final concentration: 1 μM each). Measurements were mostly done using the standard timepoints of the Seahorse equipment (3 min mixing + 2 min waiting + 3 min of measurements), except for Fig. 6A,D, where time points were 20 sec mixing + 3 min of measurements (zero time waiting), to uncover the short adrenaline effects. Cell-free wells were incubated with same medium for background correction, calculated by subtracting changes observed from the experiments in the presence of cells. OCR values obtained were normalized per amount of total protein in each well. For that, at the end of each experiment, the medium was removed, cells were PBS-washed, lysed with RIPA buffer, and total protein was quantified using the BCA Pierce protocol. Values were then normalized again by dividing the OCRs obtained by the average OCRs from the controls of each experiment.

### 4.15. Substrate oxidation test

Different substrate oxidation effects on respiration were evaluated using inhibitors for the three primary substrates that fuel mitochondria: glucose/pyruvate (12 μM UK5099, an inhibitor of the mitochondrial pyruvate carrier, MPC), glutamine (8 μM BPTES, an inhibitor of glutaminase 1, GLS-1), and long-chain fatty acids (14 μM etomoxir, an inhibitor of carnitine palmitoyl transferase 1a, CPT1a). Oxygen consumption was measured in AML-12 cells in suspension using a high-resolution O2k respirometer (Oroboros Instruments). 1 × 10^6^ cells were incubated in 2 mL of the same medium used in Seahorse experiments in the absence or presence of 2 μM thapsigargin and/or inhibitors, at 37°C with continuous stirring (400 rpm). After basal respiration, 1 μM oligomycin was injected, followed by 5 μM CCCP, to calculate maximal respiration.

### 4.16. Statistical analysis

Statistical analysis was carried out using GraphPad Prism 8 Software for at least three independent experiments (biological replicates). Student’s t test was used to compare differences between treated and untreated conditions and One-way ANOVA followed by Šidak was used to compare multiple conditions, with confidence levels set to p < 0.05. Data shown in scatter graphs are individual biological replicates, averages and standard deviations.

## Data availability

All data are contained within the manuscript.

## Acknowledgements

This work was supported mainly by the *Fundação de Amparo à Pesquisa do Estado de São Paulo* (FAPESP) under grant numbers 13/07937-8, 17/14713-0, 19/18402-4, and 21/02481-2, as well as the *Conselho Nacional de Desenvolvimento Científico e Tecnológico* (CNPq) and *Coordenação de Aperfeiçoamento de Pessoal de Nível Superior* (CAPES) line 001. We acknowledge Silvânia M. P. Neves and her animal facility crew for exceptional expert animal care, and Natália Soares Ferreira and Ricardo Bernardino de Paula from the *Centro de Facilidades de Apoio à Pesquisa* (CEFAP) for excellent technical assistance in the Confocal experiments.

## Author contributions

E.A.V.-B. and A.J.K. are responsible for concept and design; E.A.V.-B., J.V.C.-C., V.M.R. and C.C.C.S. performed experiments; E.A.V.-B., J.V.C.-C. and A.J.K. interpreted the data and discussed the results; E.A.V.-B. and A.J.K. prepared figures and drafted the manuscript; all authors revised and approved final version of manuscript.

